# Terpenoid components, antioxidant and antibacterial activities of essential oils from fresh immature and mature leaves of *Blumea balsamifera,* extracted with different hydrodistillation periods

**DOI:** 10.1101/2021.02.01.429126

**Authors:** Sirinapha Jirakitticharoen, Wudtichai Wisuitiprot, Pongphen Jitareerat, Chalermchai Wongs-Aree

**Affiliations:** Division of Postharvest Technology, School of Bioresources and Technology, King Mongkut’s University of Technology Thonburi, Bangkok, 10150, Thailand; Department of Thai Traditional Medicine, Sirindhorn College of Public Health Phitsanulok, Phitsanulok 65130, Thailand; Postharvest Technology Innovation Center, Ministry of Higher Education, Science, Research and Innovation, Bangkok 10400, Thailand

**Keywords:** *Blumea balsamifera*, leaf maturity, hydrodistillation, terpenoids

## Abstract

Volatiles and antioxidant capacities in essential oils (EOs) from fresh immature and mature leaves of *Blumea balsamifera,* extracted with different extraction periods of hydrodistillation, were investigated. There were seven major terpenoid compounds in the leaf extracts, including 2 monoterpenes of camphor and L-borneol, and 5 sesquiterpenes of silphiperfol-5-ene, 7-epi-silphiperfol-5-ene, ß-caryophyllene, ɤ-eudesmol, and α-eudesmol. Different hydrodistillation periods resulted in different quantitates and compositions of the terpenoids in EOs. The yield of EOs from the immature leaves was 1.4 times higher than the mature ones, whereas 73% of the yield was collected from the first 6 h of hydrodistillation. Camphor and L-borneol were almost collected in the first 6 h, while ß-caryophyllene, silphiperfolene, and 7-epi-silphiperfolene were above 80%, but ɤ-eudesmol and α-eudesmol were only 32 and 54% released. ß-Caryophyllene, ɤ-eudesmol, and α-eudesmol were found higher in the mature leaf EOs. Antioxidant capacities in EOs were positively related to terpenoid contents. Antibacterial activity of EOs from the immature leaves was subsequently tested. Although EOs from the hydrodistillation period of 12-18 h contained fewer terpenoid compositions, it showed the same minimum inhibitory concentration (MIC) and minimum bactericidal concentration (MBC) on *Escherichia coli* and *Pseudomonas aeruginosa,* compared to 0-6 h EOs.

## Introduction

*Blumea balsamifera* (Linn.), belonging to the Family Asteraceae, is a locally grown traditional medicine plant in Thailand. The leaves have long been used for conventional therapy in terms of healing many conditions, including skin injury, skin bruises, beriberi, eczema, dermatitis, lumbago, menorrhagia, rheumatism, and some other diseases [1, 2]. In Thailand, volatiles from *B. balsamifera* leaves have been traditionally used in heated/steamed incubation to heal women after childbirth and those who get injured. Furthermore, extracts of the leaves have been exhibited in physiological activities on free radical scavenging [3], plasmin-inhibitory [4,5], anti-obesity functions [6], antifungal activities [7], antimicrobial activities [8], anticancer [9], inhibition against the maize weevil *(Sitophilus zeamais)* [10], a reduction of blood pressure, an inhibition of sympathetic nervous system, curing over-excitement of insomnia [11].

From previous studies, 93 constituents of volatiles were reportedly found in *B. balsamifera* leaves [2], and the main components of essential oils (EOs) include L-borneol, (-)-borneol, camphor, ß-caryophyllene, ɤ-eudesmol, 10-epi-ɤ-eudesmol, ß-eudesmol, α-eudesmol [12, 13]. As a result of many bioactive compounds accumulated in *B. balsamifera* leaves, various raw materials can provide economic and social benefits. Besides, chemical differences in the composition and quantity of EOs were found in different growing regions of *B. balsamifera* [14]. In China, *B. balsamifera* plants grown in Hongshuihe village, Luodian, and Guizhou contain high L-borneol content. Moreover, the different plant organs affected the chemical composition, yield, and antioxidant activity [15]. Camphor, L-borneol, and ß-caryophyllene are among the most bioactive compounds in *B. balsamifera* leaves, giving various activities to be applied in pharmaceutical, food, cosmetic and industrial areas as antimicrobial activities, drug permeability enhancement, and anti-mutagenic effect [15–19]. Many reports studied chemical compositions of EOs in various cultivated regions [14] and in different harvesting times [20]. However, there is no report yet, studying properties of EOs from the fresh leaves to varying maturities since *B. balsamifera* is such a bush plant containing young and mature leaves on any branch. Furthermore, there was an effort to improve the extraction efficiency and prepare and separate high purity of borneol by modifying hydrodistillation and sublimation [21]. So, in the present study, volatile compositions and antioxidant capacities were investigated in hydrodistillated oils from immature and mature leaves of *B. balsamifera* using a modified hydrodistillation with different extracting times was studied. Comparison of antibacterial activities of the selected leafy EOs was then investigated.

## Materials and methods

### Plant materials

Fresh, bright green immature leaves (the 2^nd^ −4^th^ leaves from the shoot containing small soft trichomes and soft surface on upper epidermis), and dark green mature leaves (leaves containing tiny stiff trichomes and matted surface on upper epidermis) of *B. balsamifera* (Figure S1) were collected in December 2019, from 2.5 m height tree in a cultivated field at Bangkhuntian (N 13.57631; E 100.44295), Bangkok, Thailand. The plant (voucher specimen No. ttm – 0003856, Crude drug No. ttm – 1000500) was identified and certified by Mr. Nopparut Toolmal, Agricultural research officer, Thai Traditional Medicine Research Institute, Department for Development of Thai Traditional and Alternative Medicine.

### Chemical materials

Absolute ethanol was purchased from Daejung (South Korea). Trolox (6-hydroxy-2,5,7,8-tetramethyl-chromane-2-carboxylic acid), Thiophene, DPPH (2,2’-diphenyl-1-hydrazyl), ABTS (2,2’-azinobis-(3-ethylbenzothiazoline-6-sulfonic acid) diammonium salt), and Potassium persulfate were supplied by Sigma-Aldrich (USA). ß-Caryophyllene was purchased from Tokyo Chemical Industry (Japan). Camphor, and endo-borneol were provided by Alfa Aesar (USA). All chemicals and reagents were analytical grades.

### Extraction of essential oil

Fresh immature or mature leaves (500 g) were blended and then put in a 10-liter round bottom flask with deionized water (5 liters). Then, the flask was subjected to hydrodistillation using a Clevenger type apparatus (Figure S2). Hydrodistillated oil was collected every 6 h until 24 h. Each partial oil was dried over anhydrous sodium sulfate and stored in sealed vials at −20 °C until analysis.

### Determination of volatile components by GC-MS

Analysis of volatile components was followed as previously using gas chromatography-mass spectroscopy (GC-MS) [22]. An Agilent 6890N (Agilent Technologies, USA) gas chromatograph was equipped with an HP-5MS (5 % of phenyl dimethylpolysiloxane) as a fused-silica capillary column (30 m length × 0.25 mm i.d., 0.25 μm film thickness), and an Agilent 5973 Network mass selective detector (Agilent, USA). One μL of 100 ppm thiophene (v/v) (as internal standard) and 1 μL of EOs were injected. The carrier gas was Helium. The column’s temperature was 120°C, with 5 min initial hold and then increased to 250°C at a 10°C/min rate. Temperatures of injector and detector were 250°C and 200°C, respectively, manifold at 70°C with line transfer at 240°C the ionization energy was 70eV. Scan the mass spectra were in the range of 30-500 amu. Authentic standards of camphor, L-borneol, and ß-caryophyllene were used for the quantitative calculation of the compounds. The standard curves of camphor (0-2000 mg/mL; y = 0.017x – 1.6243; R^2^ = 0.9926), L-borneol (200-1000 mg/mL; y = 0.0131x – 2.1567; R^2^ = 0.9959) and ß-caryophyllene (200-800 mg/mL; y = 0.0342x – 2.9789; R^2^ = 0.9926) were prepared. Camphor, L-borneol, and ß-caryophyllene contents were expressed as microgram per one-hundred-gram fresh weight (μg /100g FW). All experiments were analyzed in triplicate.

Individual constituents of EOs were identified by comparing their Kovats retention indices (RI) relative to (C8-C30) n-alkanes on nonpolar and polar columns and comparison with literature data. Furthermore, Identification mass spectrums were matched with mass spectral library (NIST databases) or comparison with authentic compounds [23].

### Determination of antioxidant activity

#### DPPH radical scavenging assay

The total free radical scavenging capacity of the EOs was estimated according to the previously reported method [24] with slight modification using DPPH radical. The radical solution is prepared by dissolving 2.4 mg of DPPH in 100 mL of absolute ethanol. A test solution (50 μL) was added to 1.950 mL of ethanolic DPPH. The mixture was shaken vigorously and kept at room temperature for 30 min in the dark. Blank was prepared by replacing 50 μL of EOs with 50 μL of ethanol. The mixture’s absorbance was measured at 517 nm using a UV spectrophotometer (UV-1800, Shimadzu, Japan). Trolox solution was used to calibrate the standard curve at the range of 20-100 mg/mL, obtaining y = 0.0041x + 0.0262 (R^2^ = 0.9993). DPPH values were expressed as microgram of Trolox equivalent per one-hundred-gram fresh weight (μg Trolox/100g FW).

#### ABTS radical scavenging assay

The free radical scavenging activity of EOs was determined by ABTS radical cation decolorization assay [25]. ABTS cation radical was produced by the reaction between 7mM ABTS in water and 2.45 mM potassium persulfate (1:0.5) stored in the dark at room temperature for 12-16 h. ABTS solution was then diluted with ethanol to obtain an absorbance of 0.700 at 734 nm by diluting absolute ethanol before it was used. Fifty μL of *B. balsamifera* extract was mixed with 1.950 mL of ABTS radical solution. Blank was prepared by replacing 50 μL of EOs with 50 μL of ethanol. The absorbance of the mixture was spectrophotometrically measured at 734 nm. Trolox solution (20-100 mg/mL) was used to calibrate the standard curve with y = 0.006x + 0.0249 and R^2^ = 0.9996. ABTS values were expressed as microgram of Trolox equivalent per one-hundred-gram fresh weight (μg Trolox/100g FW).

### Determination of antibacterial activity

In this study, EOs from immature leaves at the hydrodistillation period of 0-6 h and 12-18 h were *in vitro* studied on antibacterial activities. The minimum inhibitory concentration (MIC) and the minimum bactericidal concentration (MBC) were measured using the broth dilution procedure as described in modified CLSI M7-A7 (2006) [26]. Three selected species from Gram-positive and Gram-negative bacteria, including *Staphylococcus aureus* (ATCC 6538), *Escherichia coli* (ATCC 8739), and *Pseudomonas aeruginosa* (ATCC 9027), were tested. The antibacterial test was certified by the Expert Centre of Innovative Herbal Products (InnoHerb) from the Thailand Institute of Scientific and Technological Research (TISTR).

Inoculum preparation of bacteria was prepared by growing each bacterial strain in media and incubating at 35 ± 2°C for 18-24 h. The standardized bacterial culture was prepared by adjusting tested bacterial suspension containing approximately 1×10^8^ CFU/mL. The stock EOs was diluted to 0.05, 0.1, and 0.5 mg/mL in a tube containing 5 mL of Mueller-Hinton broth (MHB). MIC value was observed by the presence of the turbidity of tested bacteria in the media, containing each concentration of the samples. Fifty μL of each concentration, 60% ethanol (control material), and without the sample were applied into each tube of MHB. All activity media tubes were incubated at 35 ± 2°C for 18-24 h. MIC of each EOs was determined and reported. MBC was selected from the activity media tube, which was absent of turbidity of tested bacteria in the media from MIC assay. A loopful of each selected tube was streaked on the surface of sterilized MHA plates, and the plate was incubated at 35 ± 2°C for 18 – 24 h. MBC value was observed by the presence or absence of tested bacterial growth on the MHA plates.

### Statistical analysis

All experiments were carried out in three replicates, and the data were expressed as mean values ± standard deviation of three replication of each EOs. The data was statistically analyzed by variance analysis (ANOVA) using SPSS software version 18. Mean comparisons were operated using Duncan’s multiple range test (DMRT) at P<0.05 and P<0.01.

## Results and Discussion

### Volatile components in extracts with different leaf maturities and hydrodistillation times

Hydrodistillated EOs of fresh immature and mature leaves of *B. balsamifera* afforded yellowish color. The yield of EOs from the immature leaves (501.9 mg/100g FW) was higher than that from the mature leaves (352.4 mg/100g FW). This could be caused by the higher biosynthetic rates of metabolites in young laves, resulting in increased biomass accumulation. The immature leaves raising in the positioning are intensely exposed by sunlight, which induces many secondary phytochemical compounds. Thus, the immature leaves can perceive more sunlight, affecting the efficient production of bioactive compounds in the leaves, as found in *Cistus ladanifer* [27]. The result was similar to a previous study with *B. balsamifera* in China, which studied different plant organs and other growth times [16]. Phenolics and flavonoids are a group of secondary metabolites of plants used to protect against abiotic or biotic stresses. Furthermore, secondary metabolites are from different metabolite families [28]. Thus, mature leaves could use some of the secondary metabolites used to protect themselves from the environment. Hydrodistillated EOs from the first 6 h contained the highest portion (73%) of the yields (Table 1). The longer extracting time resulted in reducing portions of the yield when the 18-24 h contained only 6.5% of the total yield. However, there was an interaction between leaf maturities and hydrodistillation times. So, EOs extracted by hydrodistillation from this plant could be collected by 6 h concerning the yield compared to the operating time.

**Table 1.** Yields of essential oils extracted from immature and mature leaves of *B. balsamifera* with different hydrodistillation periods

| Treatment |  | Weight |
| --- | --- | --- |
| Maturity | Time | (mg/100g FW) |
| Immature |  | 501.90 ± 510.48 <sup>a</sup> |
| Mature |  | 352.40 ± 497.65 <sup>b</sup> |
| F-test (Mature) |  | ** |
|  | 0-6 h | 1254.45 ± 110.53 <sup>a</sup> |
|  | 6-12 h | 222.21 ± 128.01 <sup>b</sup> |
|  | 12-18 h | 149.28 ± 88.10 <sup>c</sup> |
|  | 18-24 h | 82.64 ± 42.55 <sup>d</sup> |
| F-test (Time) |  | ** |
| (Mature x Time) |  | * |
| Immature | 0-6 h | 1333.67 ± 97.16 <sup>a</sup> |
|  | 6-12 h | 336.81 ± 31.61 <sup>c</sup> |
|  | 12-18 h | 226.01 ± 10.14 <sup>d</sup> |
|  | 18-24 h | 111.10 ± 28.78 <sup>e</sup> |
| Mature | 0-6 h | 1175.24 ± 47.73 <sup>b</sup> |
|  | 6-12 h | 107.61 ± 23.89 <sup>e</sup> |
|  | 12-18 h | 72.54 ± 40.43 <sup>e</sup> |
|  | 18-24 h | 54.19 ± 35.61 <sup>e</sup> |
| F-test |  | ** |
| C.V. (%) |  | 10.83 |

Data are expressed as mean ± SD (n = 3). Values in the same column followed by different letters indicate significant differences at $p < 0.05$ according to Duncan's multiple range test; \* = significant at $p < 0.05$ ; \*\* = significant at $p < 0.01$ level.

Seven essential volatile oils were found in the EOs from both immature and mature leaves of *B. balsamifera*, including 2 monoterpenoids of camphor, L-borneol, and 5 sesquiterpenoids of silphiperfol-5-ene, 7-epi-silphiperfol-5-ene, ß-caryophyllene, ɤ-eudesmol, and α-eudesmol (Table 2, Figure 1, Figure S3-S17). From the chemical content in Table 3, compared with thiophene, the internal standard, camphor, borneol, and 7-epi-silphiperfol-5-ene were not a significant difference between EOs from the immature and mature leaves. Since monoterpenes are biosynthesized in chloroplasts of plant cell, but camphor and borneol contents in the immature and mature leaves were not different. This evidence could bring the attention of chloroplast metabolisms between these two maturities. Although the mature leaves could contain higher chlorophylls and chloroplasts in the cells, the top of branches located of young leaves could be activated by high intense sunlight. Light was reported to activate monoterpenes synthesis in plants strongly [29] but did not affect the content of the sesquiterpene hydrocarbon. This was probably due to the different precursors of exogenous origin from photosynthesis [30]. Moreover, for hydrodistillation periods, 6 h extraction conducted the highest content of the seven selected terpenoids, compared with other extracting times. When 5 compounds of camphor, borneol, sesquiterpenoids of silphiperfolene, 7-epi-silphiperfolene, and ß-caryophyllene were above 90% eluted from the leaves at the first 6 h, y-eudesmol and α-eudesmol were extracted only 32.4 and 54.6%, respectively.

**Figure 1.**
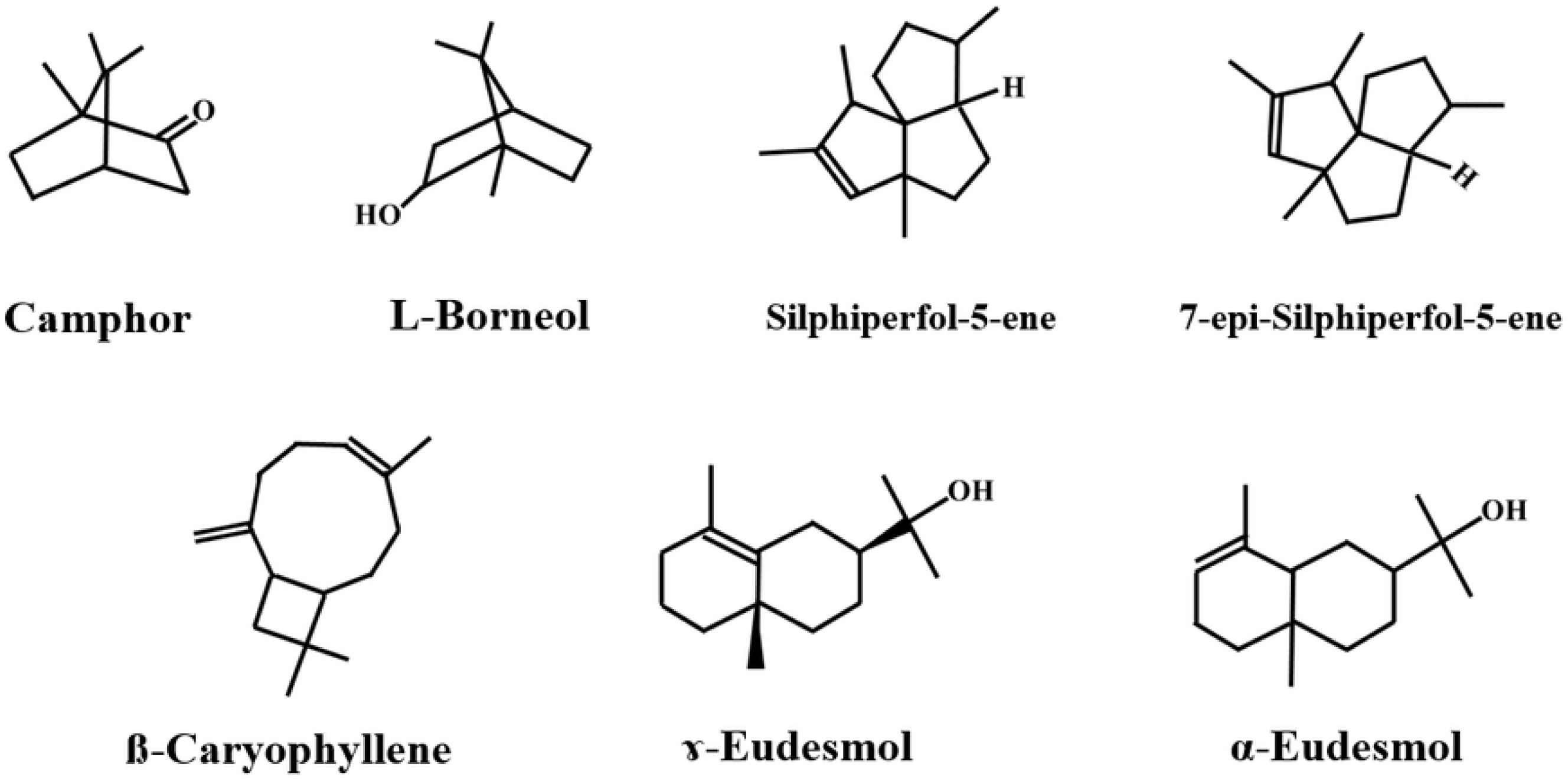
Structure of seven major terpenoids in essential oils from immature and mature leaves of *B. balsamifera,* selected from the GC-MS chromatogram

**Table 2.** Details of seven major terpenoids in essential oils from immature and mature leaves of *B. balsamifera,* selected from the GC-MS chromatogram of Figure S3-S7

| Compound name<sup>(a)</sup> | RI | MF | RT |
| --- | --- | --- | --- |
| 1.Camphor | 1121 | C<sub>10</sub>H<sub>16</sub>O | 3.540 |
| 2.L-Borneol | 1138 | C<sub>10</sub>H<sub>18</sub>O | 3.774 |
| 3.Silphiperfol-5-ene | 1403 | C<sub>15</sub>H<sub>24</sub> | 6.512 |
| 4.7-epi-Silphiperfol-5-ene | 1403 | C<sub>15</sub>H<sub>24</sub> | 6.872 |
| 5.Caryophyllene | 1494 | C<sub>15</sub>H<sub>24</sub> | 8.204 |
| 6.γ-Eudesmol | 1626 | C<sub>15</sub>H<sub>26</sub>O | 11.501 |
| 7.α-Eudesmol | 1598 | C<sub>15</sub>H<sub>26</sub>O | 11.747 |

<sup>a</sup> Compounds listed in order of elution from HP-5MS column.
RI: retention index, n-alkane
MF: molecular formula
RT: retention time

**Table 3.** Content of seven major terpenoids in essential oils from immature and mature leaves of *B. balsamifera* with different hydrodistillation periods

| Treatment |  | (g Thiophene<sup>1</sup> / 100 g FW) |  |  |  |  |  |  |
| --- | --- | --- | --- | --- | --- | --- | --- | --- |
| Maturity | Time | Camphor | L-Borneol | Silphiperfol-5-ene | 7-Epi-Silphiperfol-5-ene | β-Caryophyllene | γ-Eudesmol | α-Eudesmol |
| Immature |  | 24.55 ± 43.61<sup>a</sup> | 5.73 ± 10.74<sup>a</sup> | 1.31 ± 1.88<sup>b</sup> | 2.67 ± 1.88<sup>a</sup> | 5.70 ± 9.04<sup>a</sup> | 2.60 ± 0.80<sup>a</sup> | 1.68 ± 1.28<sup>a</sup> |
| Mature |  | 22.86 ± 39.75<sup>a</sup> | 4.00 ± 6.93<sup>a</sup> | 1.46 ± 1.94<sup>a</sup> | 2.88 ± 1.94<sup>a</sup> | 4.56 ± 6.72<sup>b</sup> | 2.03 ± 0.54<sup>b</sup> | 1.35 ± 0.95<sup>b</sup> |
| F-test (Mature) |  | ns | ns | * | ns | ** | ** | * |
|  | 0-6 h | 92.31 ± 11.38<sup>a</sup> | 18.93 ± 6.82<sup>a</sup> | 4.52 ± 0.29<sup>a</sup> | 9.11 ± 0.51<sup>a</sup> | 18.10 ± 3.13<sup>a</sup> | 3.00 ± 0.47<sup>a</sup> | 3.31 ± 0.58<sup>a</sup> |
|  | 6-12 h | 2.18 ± 0.57<sup>b</sup> | 0.46 ± 0.15<sup>b</sup> | 0.64 ± 0.18<sup>b</sup> | 1.27 ± 0.33<sup>b</sup> | 1.74 ± 0.31<sup>b</sup> | 2.59 ± 0.34<sup>b</sup> | 0.94 ± 0.09<sup>b</sup> |
|  | 12-18 h | 0.26 ± 0.11<sup>b</sup> | 0.06 ± 0.02<sup>b</sup> | 0.25 ± 0.05<sup>c</sup> | 0.47 ± 0.10<sup>c</sup> | 0.48 ± 0.10<sup>c</sup> | 2.16 ± 0.74<sup>bc</sup> | 1.08 ± 0.37<sup>b</sup> |
|  | 18-24 h | 0.07 ± 0.04<sup>b</sup> | 0.01 ± 0.01<sup>b</sup> | 0.13 ± 0.04<sup>c</sup> | 0.25 ± 0.07<sup>c</sup> | 0.20 ± 0.06<sup>c</sup> | 1.51 ± 0.29<sup>c</sup> | 0.73 ± 0.13<sup>b</sup> |
| F-test (Time) |  | ** | ** | ** | ** | ** | ** | ** |
| (Mature x Time) |  | ns | ns | ns | ns | ** | ns | ns |
| Immature | 0-6 h | 95.98 ± 15.96<sup>a</sup> | 22.40 ± 8.88<sup>a</sup> | 4.41 ± 0.31<sup>a</sup> | 9.00 ± 0.64<sup>a</sup> | 20.58 ± 2.22<sup>a</sup> | 3.39 ± 0.28<sup>a</sup> | 3.71 ± 0.56<sup>a</sup> |
|  | 6-12 h | 1.97 ± 0.58<sup>b</sup> | 0.47 ± 0.20<sup>c</sup> | 0.51 ± 0.03<sup>bc</sup> | 1.04 ± 0.07<sup>bc</sup> | 1.60 ± 0.17<sup>cd</sup> | 2.88 ± 0.17<sup>ab</sup> | 0.99 ± 0.04<sup>cd</sup> |
|  | 12-18 h | 0.22 ± 0.14<sup>b</sup> | 0.06 ± 0.03<sup>c</sup> | 0.23 ± 0.04<sup>d</sup> | 0.43 ± 0.08<sup>d</sup> | 0.47 ± 0.12<sup>cd</sup> | 2.50 ± 0.93<sup>abc</sup> | 1.23 ± 0.47<sup>c</sup> |
|  | 18-24 h | 0.04 ± 0.03<sup>b</sup> | 0.01 ± 0.00<sup>c</sup> | 0.11 ± 0.02<sup>d</sup> | 0.20 ± 0.04<sup>d</sup> | 0.16 ± 0.05<sup>d</sup> | 1.65 ± 0.32<sup>cd</sup> | 0.78 ± 0.15<sup>cd</sup> |
| Mature | 0-6 h | 88.64 ± 5.34<sup>a</sup> | 15.46 ± 1.19<sup>b</sup> | 4.64 ± 0.25<sup>a</sup> | 9.21 ± 0.44<sup>a</sup> | 15.62 ± 1.02<sup>b</sup> | 2.61 ± 0.17<sup>abc</sup> | 2.90 ± 0.15<sup>b</sup> |
|  | 6-12 h | 2.40 ± 0.59<sup>b</sup> | 0.45 ± 0.11<sup>c</sup> | 0.77 ± 0.17<sup>b</sup> | 1.50 ± 0.34<sup>b</sup> | 1.87 ± 0.39<sup>c</sup> | 2.29 ± 0.11<sup>cd</sup> | 0.90 ± 0.11<sup>cd</sup> |
|  | 12-18 h | 0.30 ± 0.08<sup>b</sup> | 0.06 ± 0.01<sup>c</sup> | 0.27 ± 0.06<sup>cd</sup> | 0.52 ± 0.10<sup>cd</sup> | 0.50 ± 0.11<sup>cd</sup> | 1.83 ± 0.43<sup>bcd</sup> | 0.94 ± 0.23<sup>cd</sup> |
|  | 18-24 h | 0.09 ± 0.04<sup>b</sup> | 0.02 ± 0.01<sup>c</sup> | 0.16 ± 0.03<sup>d</sup> | 0.29 ± 0.06<sup>d</sup> | 0.23 ± 0.05<sup>cd</sup> | 1.36 ± 0.23<sup>d</sup> | 0.67 ± 0.14<sup>d</sup> |
| F-test |  | ** | ** | ** | ** | ** | ** | ** |
| C.V. (%) |  | 25.13 | 65.12 | 11.36 | 11.04 | 17.11 | 24.99 | 18.93 |

Data are expressed as mean ± SD (n = 3) (<sup>1</sup>calculated by comparing to thiophene). Values in the same column followed by different letters indicate significant differences at p < 0.05 according to Duncan's multiple range test; ns = not significant; \* = significant at p < 0.05; \*\* = significant at p < 0.01 level.

Three main components, including camphor, borneol, and ß-caryophyllene, were further quantified with the corresponding authentic standards in Table 4. Camphor and borneol are essential ingredients used in many traditional medicines in Thailand, having therapeutic efficacies of many conventional and indigenous against diseases [31]. On the other hand, ß-caryophyllene is a sesquiterpene that has important pharmacological activities. Its exhibits a protective role of nervous system-related disorders such as depression, pain, anxiety, and Alzheimer’s disease [19]. Hydrodistillation period at 12-18 h in both immature and mature leaves obtained high amounts of ɤ-eudesmol and α-eudesmol, which are sesquiterpenoid alcohols. All eudesmol isomers (α-, ß- and ɤ-eudesmol) perform a cytotoxic effect on cancer cells [32]. Furthermore, sesquiterpenoid has a unique structure of a tricyclopentane ring, exhibiting a wide range of bioactive, biomedical, and pharmaceutical properties [33]. At the period of 6 h, EOs from the immature leaves contained quantitative of ß-caryophyllene and borneol higher than those from the mature leaves. *B. balsamifera* leaves in this study had camphor in the highest composition, followed by borneol and ß-caryophyllene (Figure 2). In contrast to some previous studies, the main components in EOs were L-borneol, followed by camphor, ß-caryophyllene, ɤ-eudesmol, isoborneol, and 1,8-cineole [10,12,13,31]. The comparison between young and mature leaves of *B. balsamifera* in China found that L-borneol, camphor, and ß-caryophyllene in the young leaves was 42.06, 1.07, and 12.24% (% relative content), whereas in mature leaves, L-borneol, camphor, and ß-caryophyllene was at 40.73, 1.12, and 11.36%, respectively [15]. Consequently, factors affecting chemical variability and yield include physiological variations, environmental conditions, and geographic variations [34].

**Figure 2.**
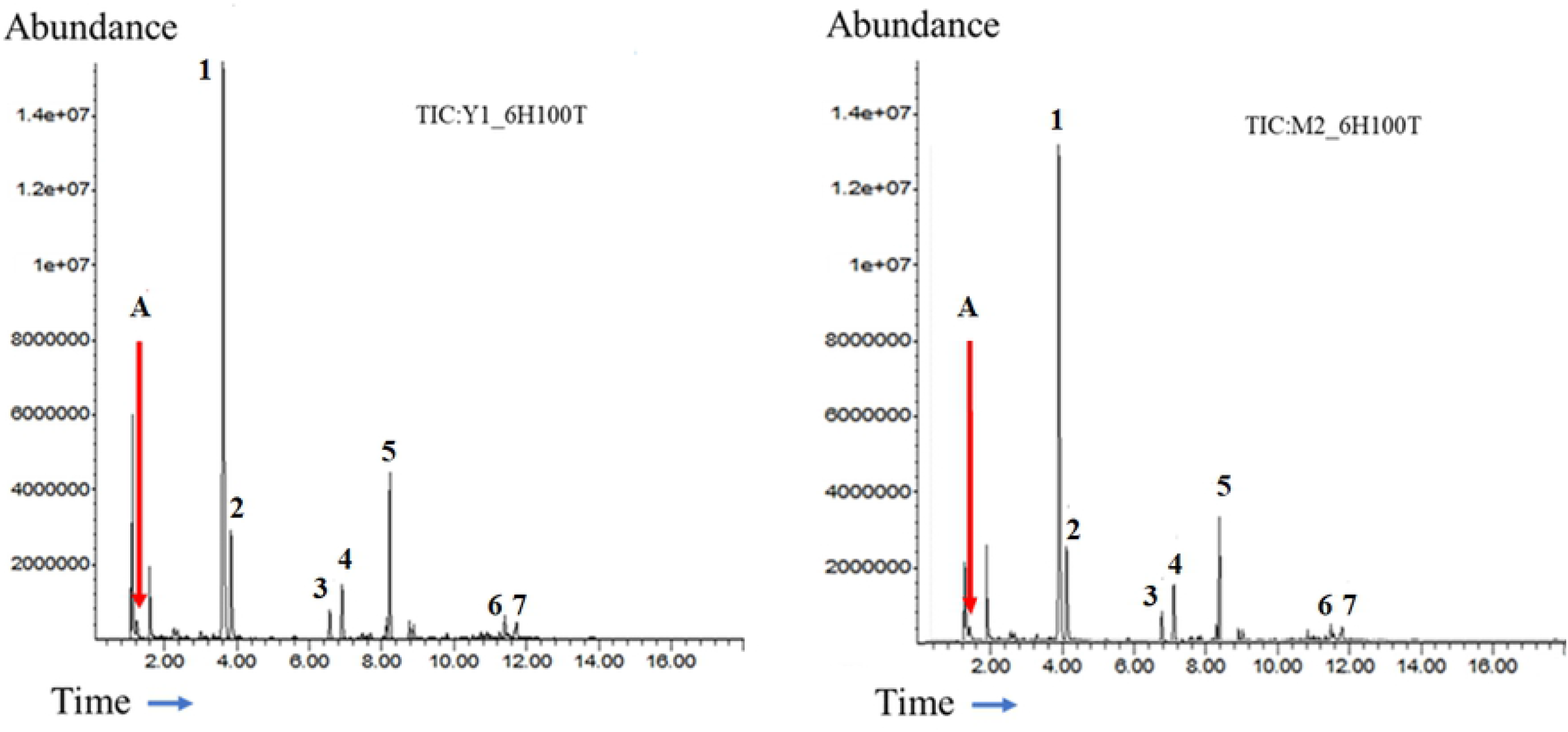
GC-MS chromatographic profile of 7 selected terpenes in essential oils (100 time ditultion) from the immature (left) and mature (right) leaves at 0-6 h hydrodistillation: A) internal standard (thiophene 100 ppm), 1) camphor, 2) L-borneol, 3) silphiperfol-5-ene, 4) 7-epi-silphiperfol-5-ene, 5) caryophyllene, 6) ɤ-eudesmol, 7) α-eudesmol

**Table 4.** Quantitative analysis of camphor, L-borneol and ß-caryophyllene in essential oils from immature and mature leaves of *B. balsamifera* with different hydrodistillation periods

| Treatment | | ($\mu\text{g}^1$ / 100 g FW) | | |
| --- | --- | --- | --- | --- |
| Maturity | Time | Camphor | L-Borneol | β-Caryophyllene |
| Immature | | 147.89 ± 261.24 <sup>a</sup> | 58.72 ± 104.95 <sup>a</sup> | 22.18 ± 35.35 <sup>a</sup> |
| Mature | | 140.12 ± 242.80 <sup>a</sup> | 45.34 ± 77.08 <sup>a</sup> | 19.02 ± 29.11 <sup>b</sup> |
| F-test (Mature) |  | ns | ns | ** |
| | 0-6 h | 559.13 ± 63.44 <sup>a</sup> | 199.40 ± 52.28 <sup>a</sup> | 73.80 ± 8.60 <sup>a</sup> |
| | 6-12 h | 14.43 ± 3.19 <sup>b</sup> | 7.12 ± 1.16 <sup>b</sup> | 6.31 ± 0.76 <sup>b</sup> |
| | 12-18 h | 1.88 ± 0.63 <sup>b</sup> | 1.15 ± 0.14 <sup>b</sup> | 1.61 ± 0.27 <sup>c</sup> |
| | 18-24 h | 0.57 ± 0.24 <sup>b</sup> | 0.44 ± 0.04 <sup>b</sup> | 0.68 ± 0.15 <sup>c</sup> |
| F-test (Time) |  | ** | ** | ** |
| (Mature x Time) |  | ns | ns | ** |
| Immature | 0-6 h | 576.11 ± 91.83 <sup>a</sup> | 225.87 ± 68.22 <sup>a</sup> | 80.48 ± 6.87 <sup>a</sup> |
| | 6-12 h | 13.35 ± 3.30 <sup>b</sup> | 7.44 ± 1.60 <sup>c</sup> | 6.06 ± 0.42 <sup>cd</sup> |
| | 12-18 h | 1.66 ± 0.79 <sup>b</sup> | 1.16 ± 0.22 <sup>c</sup> | 1.58 ± 0.31 <sup>de</sup> |
| | 18-24 h | 0.42 ± 0.16 <sup>b</sup> | 0.43 ± 0.02 <sup>c</sup> | 0.59 ± 0.12 <sup>e</sup> |
| Mature | 0-6 h | 542.16 ± 27.64 <sup>a</sup> | 172.93 ± 8.87 <sup>b</sup> | 67.11 ± 1.92 <sup>b</sup> |
| | 6-12 h | 15.50 ± 3.33 <sup>b</sup> | 6.81 ± 0.74 <sup>c</sup> | 6.55 ± 1.04 <sup>c</sup> |
| | 12-18 h | 2.11 ± 0.46 <sup>b</sup> | 1.13 ± 0.04 <sup>c</sup> | 1.64 ± 0.29 <sup>de</sup> |
| | 18-24 h | 0.72 ± 0.22 <sup>b</sup> | 0.46 ± 0.05 <sup>c</sup> | 0.76 ± 0.14 <sup>e</sup> |
| F-test |  | ** | ** | ** |
| C.V. (%) |  | 23.57 | 46.76 | 12.42 |

Data are expressed as mean ± SD (n = 3) (<sup>1</sup>calculated by comparing to the authentic standard). Values in the same column followed by different letters indicate significant differences at p < 0.05 according to Duncan's multiple range test; ns = not significant; \*\* = significant at p < 0.01 level.

### Antioxidant activity of extracts from the immature and mature leaves

The results of antioxidant capacity assays (DPPH and ABTS) were related to major terpenes from immature and mature leaves of *B. balsamifera.* Radical scavenging of DPPH and ABTS assays is related to the compound structure, which dependable on the number of active groups (OH or NH_2_) [35]. From the main composition in this study, L-borneol, ɤ-eudesmol, and α-eudesmol have hydroxyl group (OH), so that most antioxidant capacities of DPPH and ABTS could be correlated with the quantity of these 3 components. The EOs of the immature leaves comprised main components more than that of the mature leaves, and at 0-6 h of hydrodistillation, the EOs contained most compositions of the terpenes (Table 5). Extracts from the leaves at 0-6 h had the highest DPPH and ABTS activities, but the activities of EOs from the immature leaves were higher. DPPH assay has a limitation to detect only those soluble in organic solvents [36]. On the other hand, ABTS can determine both hydrophilic and lipophilic antioxidant capacity of samples [37]. So, DPPH values showed less difference in each treatment, but ABTS values have different significance between the maturity and hydrodistillation period. This implies that the photo-components extracted by the hydrodistillation method were likely to be as lipophilic higher than hydrophilic compounds.

**Table 5.**
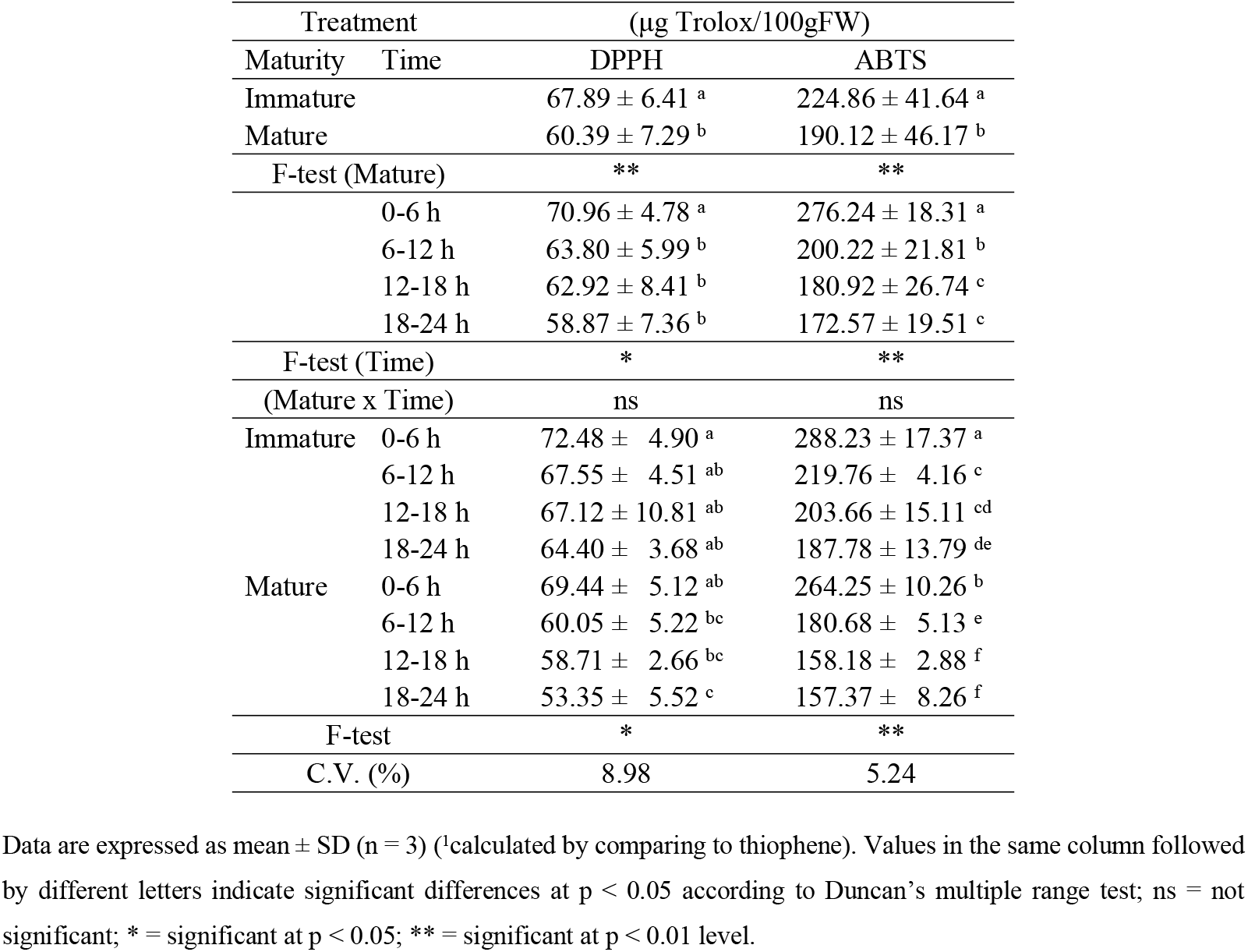
Antioxidant capacities of essential oils from immature and mature leaves of *B. balsamifera* with different hydrodistillation periods

**Table 6.** The minimum inhibitory concentration (MIC) and minimum bactericidal concentration (MBC) of essential oils from immature leaves of *B. balsamifera* extracted by hydrodistillation at the period of 0-6 h and 12-18 h against *in vitro* growth of *S. aureus, E. coli, P. aeruginosa.*

| Period Time | no. of repetitions | MICs of the extracts against the tested strains (mg/mL) |  |  | MBCs of the extracts against the tested strains (mg/mL) |  |  |
| --- | --- | --- | --- | --- | --- | --- | --- |
|  |  | *S. aureus* | *E.coli* | *P. aeruginosa* | *S. aureus* | *E.coli* | *P. aeruginosa* |
| 0-6 h | 1 | 0.5 | >5 | >5 | 1 | >5 | >5 |
|  | 2 | 0.5 | >5 | >5 | 1 | >5 | >5 |
|  | 3 | 0.5 | >5 | >5 | 1 | >5 | >5 |
| 12-18 h | 1 | 1 | >5 | >5 | >5 | >5 | >5 |
|  | 2 | 1 | >5 | >5 | >5 | >5 | >5 |
|  | 3 | 1 | >5 | >5 | >5 | >5 | >5 |

### Antibacterial activity of extracts from the immature leaves

Two different fractions of 0-6 h and 12-18 h hydrodistillated from immature *B. balsamifera* leaves were selected to evaluate *in vitro* antibacterial activity with 3 pathogenic bacteria. EOs in the 0-6 h contained high camphor, l-borneol, and ß-caryophyllene, whereas EOs in the 12-18 h had two significant elements of ɤ-eudesmol and α-eudesmol (Figure 3). The EOs were examined against a Gram-positive bacterium (*S. aureus*) and 2 Gram-negative bacteria (*E. coli and P. aeruginosa*). When *S. aureus* causes skin and soft tissue infections, bone and joint infections, bacteremia, and endocarditis [38], *E. coli* results in diarrheal disease, sepsis, and urinary tract infections [39], and *P. aeruginosa* causes gastrointestinal infection, keratitis, otitis media, and pneumonia [40–42]. Both fractions of hydrodistillation at 0-6 h and 12-18 h showed the best antibacterial effect against *S. aureus* at 0.5 mg/mL and 1 mg/mL MIC, respectively (Table 6). A previous study of EOs of *B. balsamifera* leaves from Thailand showed antimicrobial activity against *S. aureus, Bacillus cereus*, and *Candida albicans*, but there was no effect against *Salmonella enterica, Enterobacter cloacae, Klebsiella pneumoniae, E. coli*, and *P. aeruginosa* [8]. EOs of *B. balsamifera* leaves from Luodian, China had antibacterial activity against *S. aureus* and *E.coli,* whereas EOs from Hainan China performed against *S. aureus, C. albicans, Aspergillus flavus, E. coli, Listeria monocytogenes,* and *S. enterica* [14]. As a result, the EOs showed different antimicrobial activities due to various chemical compositions and different key compounds, and the extraction methods [14]. Furthermore, EOs from 0-6 h hydrodistillation conducted MBC at 1 mg/mL on *S. aureus*. Camphor is the main component of EOs, whose camphor’s aqueous solubility is more extensive than other terpenoid components in EOs [43]. Camphor’s property could probably elucidate these to penetrate through the outer membrane of bacteria [44]. The gram-negative bacteria are generally more resistant to EOs than gram-positive ones. This result could be due mainly to the outer layer of gram-negative bacteria comprised of lipopolysaccharide, which restricts hydrophobic compounds’ diffusion through its lipopolysaccharide covering [45].

**Figure 3.**
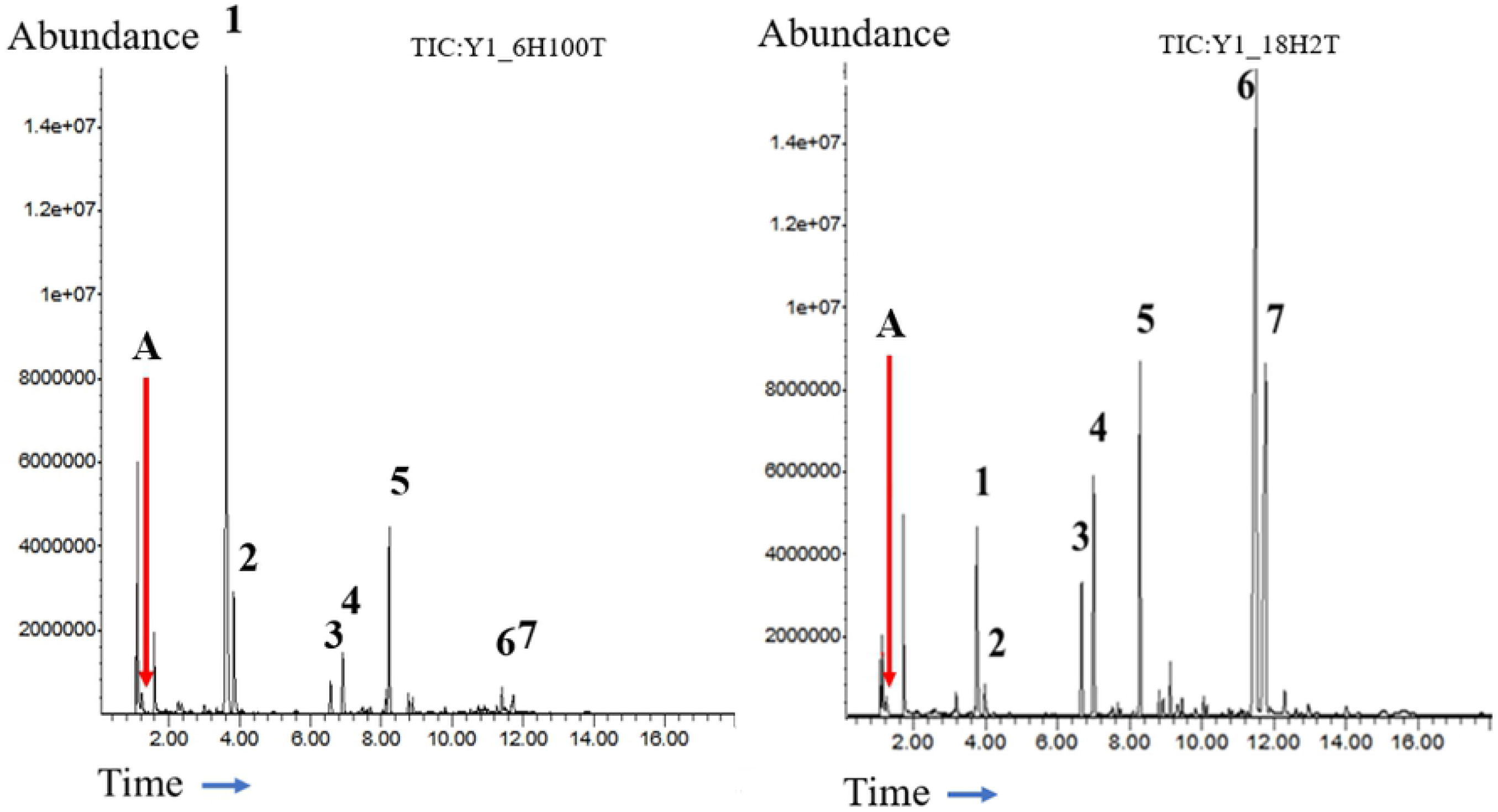
GC-MS chromatographic profile of 7 selected terpenes in essential oils from the immature leaves at 0-6 h (100 time dilution) (left) and 12-18 h (2 time dilution) (right) hydrodistillation: A) internal standard (thiophene 100 ppm), 1) camphor, 2) L-borneol, 3) silphiperfol-5-ene, 4) 7-epi-silphiperfol-5-ene, 5) caryophyllene, 6) ɤ-eudesmol, 7) α-eudesmol

## Conclusion

In this study, major of terpenoid components of EOs extracts comprised of camphor, L-borneol, silphiperfol-5-ene, 7-epi-silphiperfol-5-ene, ß-caryophyllene, ɤ-eudesmol, and α-eudesmol. EOs extracted from immature leaves contained ß-caryophyllene, ɤ-eudesmol, and α-eudesmol higher than that from mature leaves. In contrast, silphiperfol-5-ene highly accumulated in EOs from mature leaves. Contents of camphor, L-borneol, and 7-episilphiperfol-5-ene were not significantly different between immature and mature leaves. Hydrodistillation time at 0-6 h mostly extracted 7 major terpenoids from the leaves. Antioxidant capacity assays (DPPH and ABTS) were related to L-borneol, ɤ-eudesmol, and α-eudesmol. EOs extracts from immature leaves at 0-6 h of hydrodistillation period exhibited the most antioxidant activity. When the minimum inhibitory concentration (MIC) of EOs extracted from immature leaves against *S. aureus* at 0-6 h hydrodistillation period was at 0.5 mg/mL, the minimum bactericidal concentration (MBC) was at 1 mg/mL.

## Acknowledgments

This study was funded by Petchra Pra Jom Klao Ph.D. Research Scholarship (No. of agreement: 17/2560), King Mongkut’s University of Technology Thonburi. We also acknowledge the Postharvest Technology Innovation Center, Bangkok’s help in providing us some apparatus facilities.

